# Characterization of a Male Reproductive Transcriptome for *Peromyscus eremicus* (Cactus mouse)

**DOI:** 10.1101/048348

**Authors:** Lauren Kordonowy, Matthew MacManes

## Abstract

Rodents of the genus *Peromyscus* have become increasingly utilized models for investigations into adaptive biology. This genus is particularly powerful for research linking genetics with adaptive physiology and behaviors, and recent research has capitalized on the unique opportunities afforded by the ecological diversity of these rodents. However, well characterized genomic and transcriptomic data is intrinsic to explorations of the genetic architecture responsible for ecological adaptations. This study characterizes a reproductive transcriptome of male *Peromyscus eremicus* (Cactus mouse), a desert specialist with extreme physiological adaptations to water limitation. We describe a reproductive transcriptome comprising three tissues in order to expand upon existing research in this species and to facilitate further studies elucidating the genetic basis of potential desert adaptations in male reproductive physiology.

## Introduction

The rapid infusion of novel bioinformatics approaches in the fields of genomics and transcriptomics has enabled the coalescence of the fields of genetics, physiology and ecology into innovative studies for adaptation in evolutionary biology. Indeed, studies on the biology of adaptation had previously been dominated by research painstakingly documenting morphological shifts associated with ecological gradients. However, the discipline of bioinformatics has breathed new life into the field of adaptation biology. Specifically, while the morphological basis as well as the physiological mechanisms of adaptation have been explored for a variety of species in extreme environments, the genetic underpinnings of these adaptations has only recently become a larger area of research (Cheviron & Brumfield, 2011). High-throughput sequencing technology of model and non-model organisms (Ellegren, 2014) enables evolutionary biologists to conduct genome and transcriptome wide analyses and link patterns of gene selection with functional adaptations.

Studies on the genetic basis of adaptation have included a wide variety of taxa. For example, butterflies in the *Heliconius* genus have been a particularly effective study systems for determining the genetic basis of pigmentation patterns, and there is evidence of interspecific introgression for genes enabling adaptive mimicry patterns (Hines et al., 2012; The Heliconius Genome Consortium, 2012). In addition, a population genomic study in three-spine sticklebacks has elucidated many loci responsible for divergent adaptations from marine to freshwater environments (Jones et al., 2012). Another active area of adaptation genetics research focuses on species residing in extreme environments. High altitude adaptations to hemoglobin variants have been identified in multiple organisms, including humans (Lorenzo et al., 2014), several species of Andean ducks (McCracken et al., 2009a; 2009b), and deer mice, *Peromyscus maniculatus* (Storz et al., 2010; Natarajan et al., 2015). The genetic pathways responsible for physiological adaptations to desert habitats remain enigmatic; however, considerable progress has been made developing candidate gene sets for future analyses (e.g. Guillen et al., 2015; MacManes & Eisen, 2014; Marra et al., 2012; Marra, Romero, & deWoody, 2014). Functional studies will stem from this foundational research aimed at identifying the genomic underpinnings of adaptations to extreme environments; yet, it is inherently challenging and critically important to demonstrate that specific loci are functionally responsible for adaptations (Storz & Wheat 2010).

Rodents of the genus *Peromyscus* have been at the forefront of research elucidating the genetic basis for adaptation (reviewed in Bedford & Hoekstra, 2015). This diverse genus has served as an ideal platform for adaptation research spanning from the genetic basis of behavioral adaptations – such as complex burrowing in *Peromyscus polionotus* (Weber & Hoekstra 2009; Weber, Peterson & Hoekstra, 2013) – to the loci responsible for adaptive morphology – such as coat coloration in *Peromyscus polionotus leococephalus* (Hoekstra et al., 2006) and including kidney desert adaptations in *Peromyscus eremicus* (MacManes & Eisen, 2014). We are currently using *Peromyscus eremicus* as a model species for investigating the genetic bases of desert adaptations, and this paper will describe efforts to meet this research aim.

Initial steps toward understanding the genetics of adaptation must include the genomic and transcriptomic characterization of target study species (MacManes & Eisen, 2014). Toward this end, we assembled and characterized a composite transcriptome for three male reproductive tissues in the desert specialist, *P. eremicus.* This species is an exceptional example of desert adaptation, as individuals may live exclusively without water access (Veal & Clare, 2001). MacManes and Eisen (2014) assembled transcriptomes from kidney, hypothalamus, lung, and testes of this species, and they identified several candidate genes potentially underlying adaptive renal physiology. However, to our knowledge, potential physiological reproductive adaptations to water limitation have not been studied in this species or in other desert rodents. In order to pursue this novel line of adaptation research, we developed a transcriptome comprising multiple reproductive tissues in this species. Specifically, we assembled a composite reproductive transcriptome for three male reproductive tissues – the epididymis, testes, and vas deferens. Here, we describe and compare tissue specific transcriptomic data in the context of transcript abundance and relevant database searches.

## Methods

### Tissue Samples, RNA extraction, cDNA Library Preparation and Sequencing

A single reproductively mature *P. eremicus* male was sacrificed via isoflurane overdose and decapitation. This was done in accordance with University of New Hampshire Animal Care and Use Committee guidelines (protocol number 130902) and guidelines established by the American Society of Mammalogists (Sikes et al., 2011). Testes, epididymis, and vas deferens were immediately harvested (within ten minutes of euthanasia), placed in RNAlater (Ambion Life Technologies) and stored at ‐80 degree Celsius until RNA extraction. We used a standard TRIzol, chloroform protocol for total RNA extraction (Ambion Life Technologies). We evaluated the quantity and quality of the RNA product with a Qubit 2.0 Fluorometer (Invitrogen) and a Tapestation 2200 (Agilent Technologies, Palo Alto, USA).

We used a TURBO DNAse kit (Ambion) to eliminate DNA from the samples prior to the library preparation. Libraries were made with a TruSeq Stranded mRNA Sample Prep LS Kit (Illumina). Each of the three samples was labeled with a unique hexamer adapter for sequencing for identification in multiplex single lane sequencing. Following library completion, we confirmed the quality and quantity of the DNA product with the Qubit and Tapestation. We submitted the multiplexed sample of the libraries for running on a single lane at the New York Genome Center Sequencing Facility (NY, New York). Paired end sequencing reads of length 125bp were generated on an Illumina 2500 platform. Reads were parsed by tissue type according to their unique hexamer IDs in preparation for transcriptome assembly.

### Reproductive Transcriptome assembly

The composite reproductive transcriptome was assembled with reads from the testes, epididymis and vas deferens using the previously developed Oyster River Protocol for *de novo* transcriptome assembly pipeline (MacManes, 2016). Briefly, the reads were error corrected with Rcorrector v1.0.1 (Song & Florea, 2015). We used the *de novo* transcriptome assembler Trinity v2.1.1 (Haas et al., 2013; Grabherr et al., 2011). Within the Trinity platform, we ran Trimmomatic (Bolger, Lohse and Usadel, 2014) to remove the adapters, and we also trimmed at PHRED < 2, as recommended by MacManes (2014).

Next we evaluated transcriptome assembly quality and completeness using BUSCO v1.1b1 and Transrate v1.0.1. BUSCO (Simão et al., 2015) reports the number of complete, fragmented, and missing orthologs in assembled genomes, transcriptomes, or gene sets relative to compiled ortholog databases. We ran BUSCO on the assembly using the ortholog database for vertebrates, which includes 3,023 genes. The assembly was also analyzed by Transrate using the *Mus musculus* peptide database from Ensembl (downloaded 2/24/16) as a reference. The Transrate score provided a metric of *de novo* transcriptome assembly quality, and the software also generated an improved assembly comprised of highly supported contigs (Smith-Unna et al., 2015). Finally, we re-ran BUSCO on the improved assembly generated by Transrate to determine if this assembly had similar metric scores for completeness as the original assembly produced by Trinity. As alternatives to the original Trinity assembly and the optimized Transrate assembly, we proceeded with our optimization determinations by filtering out low abundance contigs from the original Trinity assembly. First we calculated the relative abundance of the transcripts with Kallisto v0.42.4 and Salmon v0.5.1. Kallisto utilizes a pseudo-alignment algorithm to map RNA-seq data reads to targets for transcript abundance quantification (Bray et al., 2015). In contrast, Salmon employs a lightweight quasi-alignment method and a high speed streaming algorithm to quantify transcripts (Patro, Duggal & Kingsford, 2015). After determining transcript abundance in both Kallisto and Salmon, we removed contigs with transcripts per million (TPM) estimates of less than 0.5 and of less than 1.0 in two separate optimization trials (as per MacManes, 2016). Finally, we evaluated these two filtered assemblies with Transrate and BUSCO to determine the relative quality and completeness of both assemblies. We chose the optimal assembly version by comparing Transrate and BUSCO metrics and also through careful consideration of total contig numbers across all filtering and optimizing versions. The chosen assembly was the Transrate optimized TPM > 0.5 filtered assembly, and this assembly was used for all subsequent analyses.

### Annotation, Transcript Abundance, and Database Searches

We used dammit v0.2.7.1 (Scott 2016) to annotate the optimized transcriptome assembly (as per MacManes, 2016). Within the dammit platform, we predicted protein coding regions for each tissue with TransDecoder v2.0.1 (Haas et al., 2013), which was used to find open reading frames (ORFs). Furthermore, dammit utilizes multiple database searches for annotating transcriptomes. These database searches include searches in Rfam v12.0 to find non-coding RNAs (Nawrocki et al., 2014), searches for protein domains in Pfam-A v29.0 (Sonnhammer, Eddy & Durbin, 1997; Finn et al., 2016), the execution of a LAST search for known proteins in the UniRef90 database (Suzek et al., 2007; Suzek et al., 2015), ortholog matches in the BUSCO databases, and orthology searches in OrthoDB (Kriventseva et al., 2015).

Next we used the assembly annotated by dammit to re-run Kallisto to determine transcript abundance within each of the three tissue types. Highly abundant transcripts were found by sorting and selecting the transcripts with the 10 highest TPM counts for each tissue. Ensembl accession numbers generated by dammit were searched within the web browser (ensembl.org) to determine the protein and gene matches corresponding to these transcripts. In addition, we used TPM counts of expression for all three tissues to generate counts of transcripts specific to and shared across tissue types.

We also downloaded the ncRNA database for *Mus musculus* from Ensembl (v 2/25/16), and we did a BLASTn (Altschul et al., 1990; Madden, 2002) search for these ncRNAs in our assembly. This database has 16,274 sequences, and we determined the number of transcript ID matches and the number of unique ncRNA sequence matches for our assembly. We also counted how many transcript matches were present in each of the tissues, and we referenced the corresponding Kallisto derived TPM values to determine the number of unique and ubiquitous transcript matches for each tissue.

We searched the annotated assembly for transporter protein matches within the Transporter Classification Database (tcdb.org). This database has 13,846 sequences representing proteins in transmembrane molecular transport systems (Saier et al., 2014). We executed a BLASTx (Altschul et al., 1990; Madden, 2002) search to find the number of transcript ID matches and the number of unique transporter protein matches within the assembly. Next we determined how many transcript ID matches were found in each of the three tissues. As previously described above, we also cross-referenced these matches with the Kallisto derived TPM values to find the number of ubiquitous and unique transcript matches by tissue type.

Of note, the code for performing all of the above analyses can be found at https://github.com/macmanes-lab/peer_reproductive-transcriptome/blob/master/code.md. The data files that are in Dropbox, which will later be submitted to Dryad, can be found at https://www.dropbox.com/home/PEReproductiveTranscriptome/To%20Submit%20to%20Dryad.

## Results and Discussion

### Reproductive Transcriptome assembly

There were 45-94 million paired reads produced for each of the three transcriptome datasets, yielding a total of 415,960,428 reads. The raw reads are available at the European Nucleotide Archive under study accession number PRJEB13364.

We assembled a *de novo* composite reproductive transcriptome with reads from testes, epididymis and vas deferens. The evaluation of alternative optimized assemblies allowed us to generate a substantially complete transcriptome of high quality. The alternative assemblies had raw Transrate scores ranging from 0.194-0.208 (**Table 1)**. However, the scores for the improved assemblies generated by Transrate, consisting of only highly supported contigs, ranged between 0.295-0.349, which is well above the threshold Transrate score of 0.22 for an acceptable assembly. The BUSCO results indicated that the assemblies were highly complete, with complete matches ranging from 73-90% of vertebrate orthologs (**Table 2**). These BUSCO benchmark values are consistent with the most complete reported assessments for transcriptomes from other vertebrate taxa (busco.ezlab.org). Furthermore, our BUSCO values exceed that of the only available reported male reproductive tissue (from a coelacanth: *Latimeria menadoensis* testes), which was 71% complete (Simão et al., 2015). The assembly version which was of highest quality in relation to the Transrate metrics was the Transrate optimized Trinity assembly; specifically, the optimized Transrate score was 0.3495, and the percent coverage of the reference assembly was also highest, with 45% of the mouse database represented. This assembly was highly competitive for completeness, as indicated by the BUSCO metric of 85% orthologs found. However, this assembly had an exorbitantly high number of contigs (657,952 contigs), which is nearly an order of magnitude more contigs than the next best performing assembly: the Transrate optimized TPM > 0.5 filtered assembly (78,424 contigs). In consideration of the dramatically more realistic contig number for the Transrate optimized TPM > 0.5 filtered assembly, and in light of its second best performance for Transrate score (0.3013), reasonable Transrate mouse reference assembly coverage (37%), and sufficiently high BUSCO completeness (73% orthologs found), we chose this assembly as our optimized transcriptome. Therefore, we proceeded with this optimized assembly version as our finalized transcriptome assembly for our analyses, which is available in Dropbox (to be posted on Dryad after this manuscript’s acceptance).

**Table 1:** Transrate results for the reproductive transcriptome assembly produced by different optimization methods. * This is the score of the Transrate optimized assembly in Table 2.

| Assembly | Transrate Score | Optimized Score* | # Read Pairs (fragments) | Contigs (n_seqs) | # Good Contigs | %Good Contigs |
| --- | --- | --- | --- | --- | --- | --- |
| Trinity Original | 0.1944 | 0.3492 | 207,980,214 | 856,711 | 657,952 | 0.77 |
| Filter TPM<0.5 | 0.1672 | 0.3013 | 207,980,214 | 147,966 | 78,424 | 0.53 |
| Filter TPM<1.0 | 0.156 | 0.2854 | 207,980,214 | 80,165 | 54,140 | 0.68 |

**Table 2:** BUSCO metrics for the reproductive transcriptome assembly produced by different optimization methods.

| Assembly | % Complete | %Duplicated | %Fragmented | %Missing |
| --- | --- | --- | --- | --- |
| Trinity original | 90 | 49 | 3.4 | 5.5 |
| Transrate Optimized | 85 | 44 | 4.3 | 9.7 |
| Filter TPM<0.5 | 85 | 38 | 3.0 | 11 |
| Transrate TPM<0.5 | 73 | 31 | 3.9 | 22 |
| Filter TPM<1.0 | 80 | 28 | 2.8 | 16 |
| Transrate TPM<1.0 | 74 | 25 | 3.4 | 21 |

### Annotation, Transcript Abundance and Database Searches

The reproductive transcriptome assembly annotations were produced by dammit, and they are available through Dropbox (this file, and all other data files will be posted on Dryad after acceptance) in a gff3 file format. Furthermore, TransDecoder was used to predict coding regions in the assembly. TransDecoder predicted that 49.5% (38,342) of the transcripts (78,424 total) contained ORFs, of which 63.9% (24,808) had complete ORFs containing a start and stop codon. The predicted protein coding regions generated by TransDecoder are reported in five file types, and they are available on Dropbox. Furthermore, the Pfam results yielded 30.7% of transcripts (24,107) matching to the protein family database. In contrast, the LAST search found that 75.9% of transcripts (59,503) matched to the UniRef90 database. We have uploaded the homology search results generated by Pfam and UniRef90 matches onto Dropbox. In addition, 1.04% (816) of transcripts matched to the Rfam database for ncRNAs, and these results are posted in Dropbox. Of note, 80.1% (62,835) of the transcripts were annotated using one of more of the above described methods (the dammit.gff3 file is posted in Dropbox), and it is this final annotated assembly that was used for all subsequent analyses.

The transcripts with the 10 highest TPM counts generated by Kallisto for each tissue type corresponded with protein matches in Ensembl (**Tables 3-5**). All three tissues had highly abundant transcripts for mitochondrially encoded cytochrome c oxidase subunits, which are involved in cellular respiration. Highly abundant testes proteins included protamine 2 – which is involved in spermatogenesis – and sperm autoantigenic protein 17 – a zona pellucida binding protein. Similarly, the highly abundant epididymis transcripts consisted of a protein involved in spermatozoa maturation, cysteine-rich secretory protein 1, as well as Cd52 (also known as epididymal secretory protein E5). In contrast, the vas deferens had abundant transcripts for proteins involved in muscle contraction, specifically several actin subunits. The highly abundant transcripts in all three tissues were consistent with our expectations of their physiology and in keeping with findings in humans. Specifically, protamine-2 was the most highly expressed gene in the human testes, and zona pellucida binding proteins were also highly expressed in human testes (Djureinovic et al., 2014).

**Table 3:** Testes transcript annotations for top 10 TPM results generated by Kallisto.

| Transcript ID | Ensembl Accession Number | TPM | Protein | Gene |
| --- | --- | --- | --- | --- |
| Transcript_72932 | ENSRNOP00000014866 | 23136.8 | Sperm autoantigenic protein 17 | Spa17 |
| Transcript_56114 | ENSMUSP00000047925 | 19154.7 | Protamine 2 | Prm2 |
| Transcript_37923 | ENSMODP00000000682 | 18016.4 | Uncharacterized Protein | N/A |
| Transcript_197 | ENSLACP00000009213 | 4841.98 | ubiquitin A-52 residue ribosomal protein fusion product 1 | Uba52 |
| Transcript_56089 | ENSMUSP00000080993 | 4236.73 | mitochondrially encoded cytochrome c oxidase I | Mt-co1 |
| Transcript_21368 | ENSRNOP00000065673 | 4186.17 | N/A | N/A |
| Transcript_70782 | N/A | 4085.53 | N/A | N/A |
| Transcript_67718 | ENSMUSP00000080997 | 3840.31 | mitochondrially encoded cytochrome c oxidase III | Mt-co3 |
| Transcript_51810 | N/A | 3013.74 | N/A | N/A |
| Transcript_67719 | ENSRNOP00000046414 | 2950.84 | mitochondrially encoded cytochrome c oxidase II | Mt-co2 |

**Table 4:** Epididymis transcript annotations for top 10 TPM results generated by Kallisto.

| Transcript ID | Ensembl Accession Number | TPM | Protein | Gene |
| --- | --- | --- | --- | --- |
| Transcript_48473 | N/A | 99593 | N/A | N/A |
| Transcript_48471 | ENSMUSP00000026498 | 59217.6 | cysteine-rich secretory protein 1 | Crisp1 |
| Transcript_43156 | N/A | 26929.8 | N/A | N/A |
| Transcript_56089 | ENSMUSP00000080993 | 24654.4 | mitochondrially encoded cytochrome c oxidase I | Mt-co1 |
| Transcript_67718 | ENSMUSP00000080997 | 19450.7 | mitochondrially encoded cytochrome c oxidase III | Mt-co3 |
| Transcript_67719 | ENSRNOP00000046414 | 16564.4 | mitochondrially encoded cytochrome c oxidase II | Mt-co2 |
| Transcript_65120 | ENSRNOP00000020688 | 14990.1 | CD52 molecule | Cd52 |
| Transcript_48472 | ENSMUSP00000026498 | 12385.4 | cysteine-rich secretory protein 1 | Crisp1 |
| Transcript_49280 | ENSMUSP00000026498 | 10974.7 | cysteine-rich secretory protein 1 | Crisp1 |
| Transcript_73235 | ENSSTOP00000007662 | 9807.83 | Fatty acid-binding protein, adipocyte | Fabp4 |

**Table 5:**
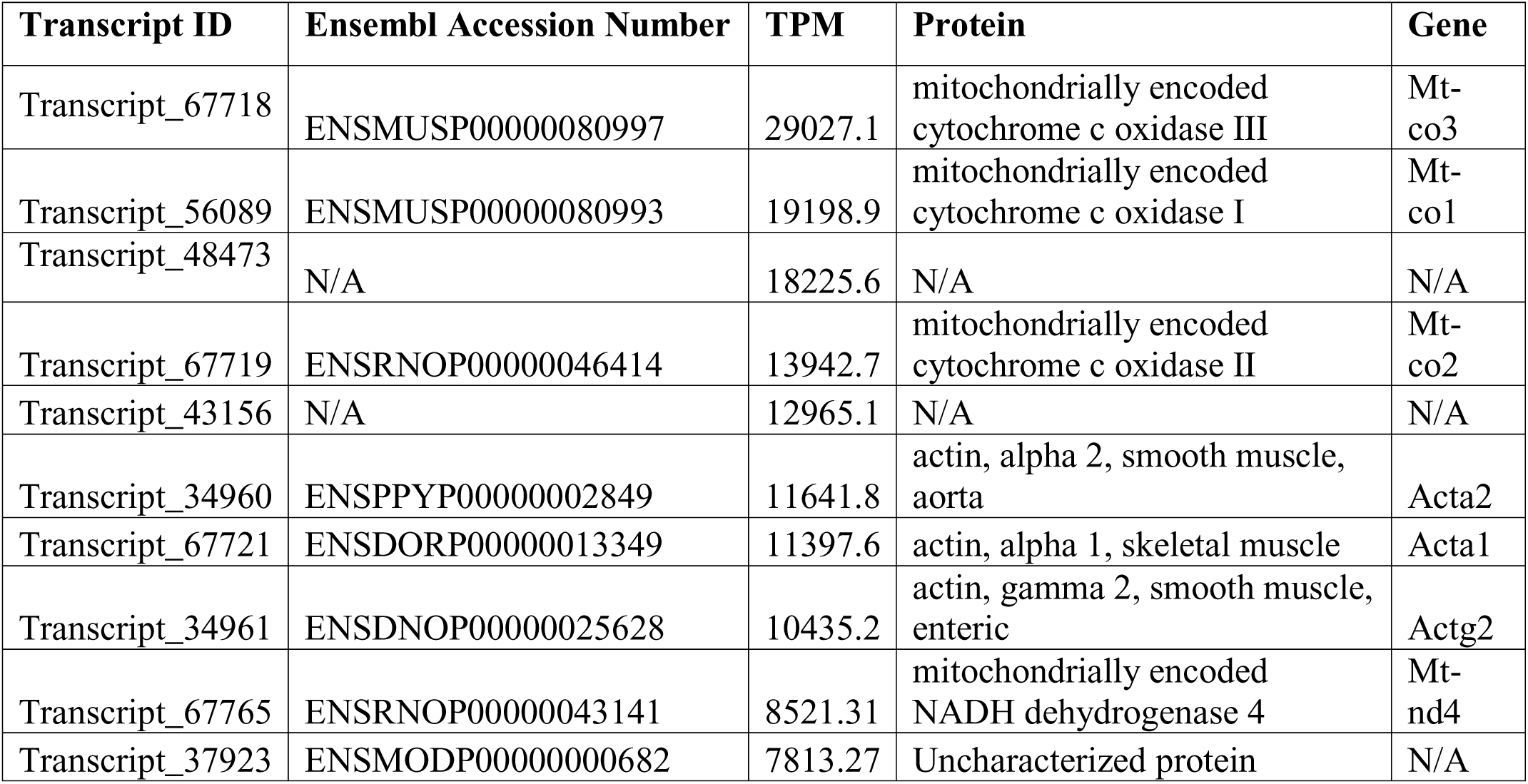
Vas Deferens transcript annotations for top 10 TPM results generated by Kallisto.

The Kallisto generated TPM counts of expression (available on Dropbox) were also utilized to determine which transcripts were ubiquitous and specific to the three tissue types, which we have depicted in a Venn diagram format (**Figure 1**). The assembly consisted of 78,424 different transcript IDs, of which 64,553 were shared across all three tissues. The number of unique transcripts were as follows: 3,563 in testes, 342 in epididymis, and 502 in vas deferens. The relatively large number of unique transcripts in the testes is consistent with previous findings which describe the testes as the tissue with the highest number of tissue-enriched genes in the human body (Uhlen et al., 2015).

**Figure 1:**
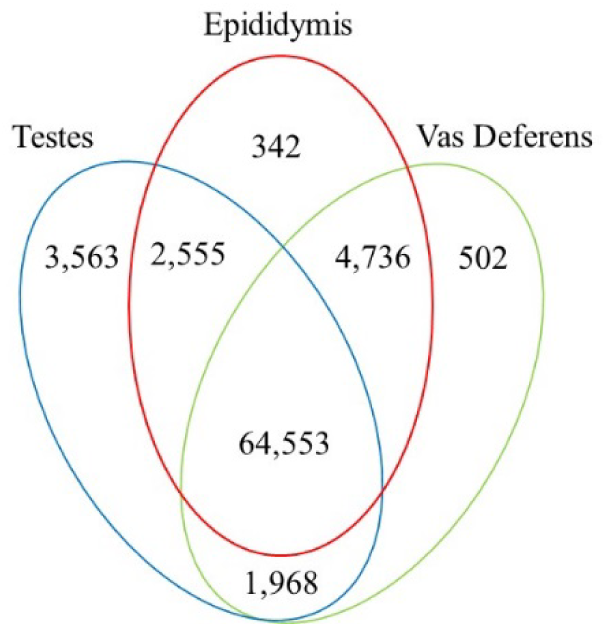
Venn Diagram of transcript expression differences and similarities between the three reproductive tissues. The total number of transcripts is 78,424.

In addition, we searched for *Mus musculus* ncRNA sequence matches within our assembly. There were 15,964 transcript matches, which correspond to 2,320 unique ncRNA matches, and they are posted on Dropbox. The transcript matches by tissue type were found using the Kallisto TPM determinations, and they were as follows, testes: 15,260, epididymis: 15,552, and vas deferens: 15,558. A Venn Diagram depicts unique and shared transcript matches by tissue type (**Figure 2**). The majority of transcript matches were ubiquitous to all three tissues (14,724), and there were far fewer tissue specific matches. The testes had more unique transcript matches (185) than the epididymis (26) or the vas deferens (45). These findings are consistent with our results above regarding the relative numbers of total unique transcripts in the assembly by tissue type.

**Figure 2:**
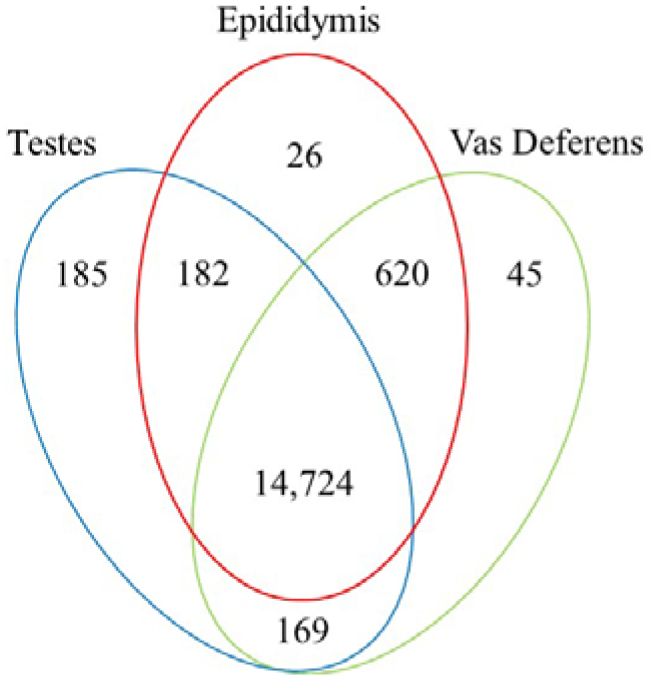
Venn Diagram of transcript matches between the three reproductive tissues to ncRNA sequences in *Mus musculus.* The total number of transcript matches across the tissue types is 15,964.

Our search for transporter protein matches within the Transporter Classification Database yielded 7,521 different transcript matches, corresponding to 1,373 unique transporter protein matches, and they are posted on Dropbox. The number of transcript matches was highly similar between the tissue types (testes 7,025; epididymis 7,115; vas deferens: 7,071). We generated a Venn Diagram to display the numbers of shared and unique transcript matches to the transporter protein sequences (**Figure 3**). Most transcript matches were present in all three tissues (6,472), and there were relatively few unique matches in the three tissue types. However, the testes had the highest number of unique transcript matches (215) relative to the epididymis (19) and the vas deferens (37). These results are in keeping with those reported for the ncRNA sequence matches and the complete assembly dataset. Furthermore, our BLASTx search of this transporter protein database yielded transcript matches for multiple solute carrier proteins. We are particularly interested in solute carrier proteins because previous research has found candidate genes in this protein family for desert adaptations in kidneys of the kangaroo rat (Marra et al. 2012; Marra, Romero & deWoody 2014) and the Cactus mouse (MacManes and Eisen 2014). In addition, we had multiple matches to aquaporins, which are water channels allowing transport across cellular membranes. One transcript matched specifically to Aquaporin 3, a sperm water channel found in mice and humans, which is essential to maintaining sperm cellular integrity in response to the hypotonic environment within the female reproductive tract (Chen et al., 2011).

**Figure 3:**
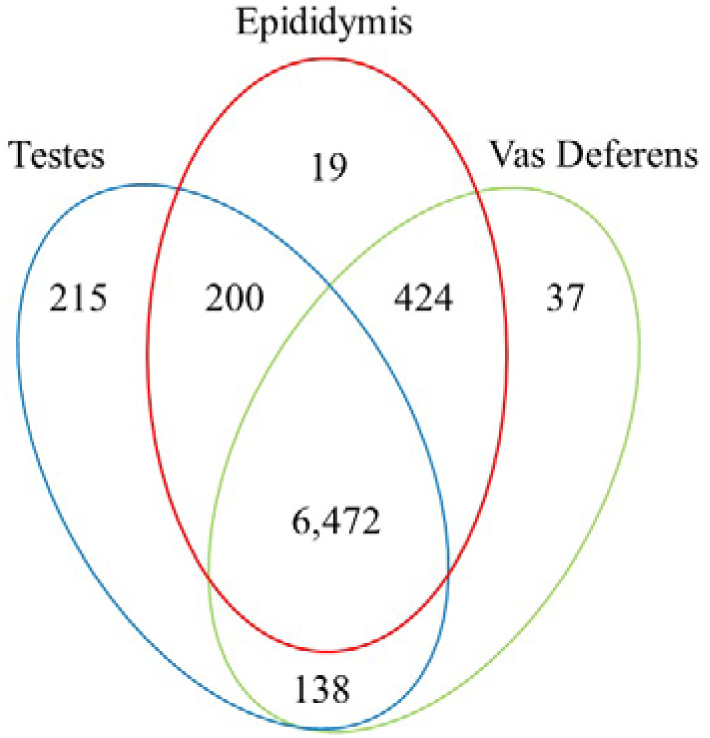
Venn Diagram of transcript matches between the three reproductive tissues to protein sequences in the Transporter Classification Database. The total number of transcript matches across the tissue types is 7,521.

## Conclusions

This study describes a composite transcriptome from three male reproductive tissues in the desert specialist *Peromyscus eremicus*. Our analyses include quality and completeness assessments of this reproductive assembly, which was generated using reads from testes, epididymis and vas deferens of a male Cactus mouse. We also describe transcript expression levels, generate annotations, and search relevant databases for ncRNAs and transporter protein sequences. Finally, we describe the degree of ubiquity between transcripts among the three tissues as well as identify transcripts unique to those tissues. Our future research will investigate potential male reproductive physiology adaptations to water limitation in Cactus mouse, and the characterization of this reproductive transcriptome will form the foundation of studies along this vein. Moreover, this research contributes transcriptomic materials to a larger body of work in the expanding field of adaptation genetics, which benefits tremendously from enhanced opportunities for comparative analyses.

## Dropbox Data File List

Final Annotated Reproductive Tissue Transcriptome: *reproductive.annotated.fasta* (127 MB)

Transdecoder (Five Files):

*transdecoder.gfí3* (49 MB)
*transdecoder.pep* (27 MB)
*transdecoder.cds* (62 MB)
*transdecoder.mRNA* (172 MB)
*transdecoder.bed* (10 MB)

Pfam Annotation: *reproductive.pfam.gffô* (32 MB)

Rfam Annotation: *reproductive.rfam.gffô* (191 KB)

Dammit Annotation: *reproductive.dammit.gff3* (98 MB)

UniRef90 Annotation: *reproductive.uniref.gffô* (9.5 MB)

Kallisto Results for Annotated Transcriptome (Three Files):

*kallisto.testes. tsv* (3.4 MB)
*kallisto.epi.tsv* (3.4 MB)
*kallisto.vas.tsv* (3.4 MB)

ncRNA Database Matches (three files):

*epi.tpm.plus.ncRNA.txt* (4.3 MB)
*vas.tpm.plus.ncRNA.txt* (4.3 MB)
*testes.tpm.plus.ncRNA.txt* (4.3 MB)

tcdb Datatbase Matches (three files):

*epi.tpm.plus.tcdb.txt* (946 KB)
*vas.tpm.plus.tcdb.xt* (946 KB)
*testes.tpm.plus.tcdb.xt* (946 KB)

